# An integrated approach to assess Knowledge, Attitude and Practices (KAP) regarding major Neglected Tropical Diseases endemic in the Mbengwi health district (North West Region, Cameroon)

**DOI:** 10.1101/658849

**Authors:** Laurentine Sumo, Cédric G Lenou-Nanga, Ngum H Ntonifor, Nicanor Chenkumo-Kengmoni, Vanessa T Amana-Bokagne, Chembo G Awah, Yannick Niamsi-Emalio, Hugues C Nana-Djeunga

## Abstract

**Background:** Preventive chemotherapy (PCT) is the main strategy currently used to control and/or eliminate onchocerciasis (Oncho), lymphatic filariasis (LF) and Soil Transmitted Helminthiasis (STH), and community participation (through implementation of MDA or adherence to PCT) is critical to achieve this goal. However, these Neglected Tropical Diseases (NTDs) are still persisting in most endemic areas as a consequence of sub-optimal treatment coverage, the presence of systematic non-compliers in communities … This study aimed at investigating whether the knowledge, attitudes and practices of populations about these NTDs can explain the poor trends towards elimination.

**Methodology:** A cross-sectional survey was carried out in the Mbengwi Health District (North West Region, Cameroon) using the cluster sampling technique. Clusters were selected using the Probability Proportionate to Estimate Size strategy. In each cluster, the random walk technique was used for the selection of households, and a structure questionnaire was administered to 2-3 of its members.

**Principal Findings:** A total of 254 households from 26 clusters were visited, and 514 individuals were interviewed. The sex ratio of interviewees (1.08) was unbiased, and their ages ranged between 10 and 99 years old. Though most of the respondents declared having already heard of these NTDs (41.2%, 73.7% and 89.9% for Oncho, LF and STH respectively), only a minority of them were aware of correct response of how they are acquired/transmitted (3.7%, 6.8% and 12.5% for Oncho, LF and STH respectively), and prevented (23.1%, 18.9% and 47.2% for Oncho, LF and STH respectively). Even when respondents were aware that medicines were useful to prevent and/or treat these NTDs, almost none of them knew the drug used or the treatment frequency.

**Conclusion/Significance:** This study reveals that interviewees exhibit poor knowledge, attitudes and practices as regards to these NTDs, although they are endemic in the study area and PCTs given yearly since a while. These misconceptions can seriously affect the adherence and contribution of populations to the success of PCTs, and it appears compulsory to improve individual knowledge, with a focus on the importance and rationale behind MDA, to optimize their attitudes and practices, especially community participation to PCTs.

**Author summary:** The control and/or elimination of Neglected Tropical Diseases (NTDs) are currently on top of the agenda of endemic countries’ control programs and stakeholders. Ivermectin- and Albendazole/Mebendazole-based mass distribution is currently the main strategy to control/interrupt transmission of onchocerciasis, lymphatic filariasis and soil-transmitted helminthiasis, and adherence of communities is compulsory for the success of this approach. Despite the success registered in the fight against these diseases, the trend towards elimination remains unsatisfactory in many endemic areas. This study was carried out to assess whether the perceptions, attitudes, and practices of the Mbengwi health district (North West Region, Cameroon) populations regarding these three parasitic diseases can explain the poor trend towards elimination. A cross-sectional study revealed important misconceptions about these NTDs amongst most of the respondents, which can clearly affect their adherence and contribution to the success of preventive chemotherapies, and seriously slower the momentum towards elimination.

## Introduction

Neglected Tropical Diseases (NTDs) are among the world’s most common conditions prevailing in tropical and subtropical settings and disproportionately affecting the poorest [1]. Since the update of the NTD portfolio in 2017 by the addition of chromoblastomycosis and other deep mycoses, scabies and other ectoparasites and snakebite envenoming [2], the burden of NTDs is yet to be updated. The previously known 17 NTDs were recognized to affect more than 2 billion people in developing countries in 2013, STH infections (ascariasis, trichuriasis, and ancylostomiasis/necatoriasis) accounting for more than three-quarters of the total NTD infections prevalence. In addition to their morbidity, the number of death attributable to the 17 NTDs prioritized by the WHO plus “other NTDs” in 2013 was estimated to be equivalent to more than half of the malaria deaths and more than double due to tuberculosis [3–5].

Despite their high burdens, these NTDs are preventable and/or treatable. The control and/or elimination of these NTDs are now on top of the agenda of endemic countries’ control programs and stakeholders in their efforts to achieve Millennium Development Goals for sustainable poverty reduction [3, 6]. Five public-health interventions – (i) preventive chemotherapy (PCT), (ii) innovative and intensified disease management, (iii) vector control and pesticide management, (iv) safe drinking-water, basic sanitation and hygiene services, and education, (v) zoonotic disease management – have been identified to accelerate the prevention, control, elimination and eradication of NTDs, more effective impact being achieved when these interventions are combined [1].

Success in controlling these NTDs have recently been achieved in a number of countries [7], but the trends towards elimination remains poor or unsatisfactory. Indeed, these NTDs are still persisting in most endemic areas, especially when long-term sustainable efforts are required as it is the case of the five major ‘tool-ready’ NTDs (lymphatic filariasis, onchocerciasis, schistosomiasis, soil-transmitted helminthiasis and trachoma). Among the reasons identified to hinder the elimination of these NTDs, sub-optimal treatment coverage and the presence of systematic non-compliers in communities are significant. The implication and contribution/participation of populations in this machinery, through implementation of MDA or adherence to PCT, are highly needed and can appear critical in the momentum towards elimination. For instance, it was demonstrated that black fly biting rates have declined by 89-99% after members of some communities in Northern Uganda have been mobilized to "slash and clear" the breeding sites of the vegetation that represents the primary onchocerciasis vector larvae attachment point [8]. It was argued that water, sanitation, and hygiene interventions as well as their combination, are effective at reducing STH infections [9], though most of the children interviewed in the framework of a study conducted in Senegal declared that they usually defecate somewhere else, though their communities were well endowed with pit latrines [10]. This means that the appropriation of control measures by the populations is a key point, and this can be made through appropriate education of endemic communities to improve their knowledge regarding these infections.

The objective of this study was to assess knowledge and perceptions of populations, in relation to their attitudes and practices, regarding the most prevalent NTDs in the Mbengwi Health District (North West Region, Cameroon).

## Methods

### Ethical statement

Ethical clearance was granted by the Institutional Review Board of the Faculty of Science of the University of Bamenda. After approval of the local administrative and traditional authorities, the objectives and schedules of the study were explained to community leaders and to all eligible individuals selected in the household. Verbal agreements were obtained from those who agree to participate. The approval of parents or legal guardians of minors was necessary before any procedure. An individual code was attributed to each participant for anonymous data analysis.

### Study area and populations

Mbengwi (6°01’N, 10°00’E) is the capital of the Momo Division (North West Region, Cameroon), situated at about 20 km to the west of Bamenda town, the Regional capital, and at an altitude ranging from 900m to 2000m above sea level. The hydrographic network is intricate with many streams and springs (the Momo division is mainly irrigated by the Momo River), and the area has a huge hydroelectricity potential afforded by the Abi falls. The climate of the Mbengwi health district is of mountain monsoon equatorial type, with a short dry season running from September to March, and a long rainy season extending from March to September. The annual average rainfall is 2022.3 mm, and the average maximum yearly temperature is 30°C [11–12]. This area is characterized with a long valley stretch surrounded by mountains (most of them counting at least 1500m in height) and lies in the transitional zone between the western grass fields and forest region. It is highly dominated by the savannah vegetation (especially on the hills) which favors animal rearing. The valley is mainly made up of trees (palm trees, raffia palms and many fruit trees), and is favorable for settlement and agriculture. Forest characteristics are highly observed at the western part of the area and favors the cultivation of cash crops like cocoa [11–12].

### Study design and sampling strategy

This study was designed as a representative 2-staged clustered cross-sectional household and questionnaire-based survey. Clusters (communities) were selected using the Probability Proportional to Estimate Size (PPES) strategy [13], and in each selected community, the random walk technique [14] was used for households’ selection. In each selected household, 1-3 individuals from different age classes – 10-20 years old, 21-30 years old and 31 years and over (usually the head of household) – were randomly selected and underwent in-depth interviews. A structured questionnaire was administered by three trained interviewers, assisted by autochthones speaking the local language, to collect participants’ socio-demographic and socio-economic characteristics (gender, age, village of residence, profession, educational level, means of locomotion), and to assess whether they have ever heard about the targeted NTDs (onchocerciasis, lymphatic filariasis and soil transmitted helminthiasis), as well as the level of their knowledge in relation to their attitudes and practices as regards to the mode of transmission, clinical manifestations and the means to prevent and treat these parasitic diseases.

### Data analysis

All relevant data were recorded into a purpose-built Microsoft Access database and subsequently transferred to R software, version 3.4.4 (The R Foundation for Statistical Computing) for statistical analysis. Categorical variables (gender, occupation, urbanization) were summarized using frequencies, and continuous variables (age) were described using median (interquartile range, IQR).

To better evaluate respondents’ knowledge, attitude and practices, a score between 0 and 2.5 was allocated to each question, either related to participants’ knowledge or to their attitudes and practices as regards to each targeted NTDs (Supplementary material Table S1). The overall score for a given study participant was the sum of all responses for a specific disease. Respondents whose scores were equal and above the mean (over a total of 10) were considered as having ‘good knowledge, attitude and practice’ (coded as 1), while those below the mean were considered as having ‘poor knowledge, attitude and practice’ (coded as 0). Associations between a KAP score and different respondents’ socio-demographic and socio-economic characteristics were tested using the Chi-Square and Fisher exact probability tests.

The Classification and Regression Tree (CART) was used to assess the association between socio-demographic characteristics and respondents’ knowledge, attitude and practice scores for each of the targeted diseases. Indeed, CART is a non-parametric multiple regression approach that both avoids multicollinearity issues and explains a categorical dependent variable by defining groups of subjects with similar behaviors [15–16], while taking into account all interactions between different covariates [17]. CART then evaluates all the possible thresholds and separates the dependent variable into two groups, the procedure being repeated recursively until an optimal criterion is obtained [18]. This leads to the selection and editing of an optimal decision tree where the leaves correspond to similar behavior classes. Odds ratio (OR) with 95% CI generated using logistic regression models were then used to describe the strength of association between the response variable or outcome (KAP scores) and independent variables (behavioral classes) before and after controlling for possible confounding variables. Non-overlapping 95% CI or *p*-values ≤5% were considered as statistically significant.

## Results

### Socio-demographic and socio-economic status of study participants

Table 1 summarizes socio-demographic and socio-economic characteristics of the survey respondents. A total of 254 households selected in 26 clusters were visited across the Mbengwi health district, and 514 participants interviewed. The sex ratio (Male / Female) of enrollees was 1.08, their age ranging between 10 and 99 years old (median: 35 years old; IQR: 23 – 49 years old). The educational level of most of the study participants was primary/secondary (84.1%), only a few being illiterate. More than half (50.4%) of the interviewees were farmers, living almost exclusively in rural (46.5%) and semi-rural/semi-urban (40.8%) settings, most of them using motorbike (58.2%) as means of locomotion.

**Table 1.** Socio-demographic and socio-economic characteristics

| Characteristics | Number (%) | Characteristics | Number (%) |
| --- | --- | --- | --- |
| **Urbanization** |  | **Sex** |  |
| Rural | 230 (46.5) | Female | 247 (48) |
| Semi urban | 202 (40.8) | Male | 267 (52) |
| Urban | 63 (12.7) | **Age** |  |
|  |  | [10 – 21[ | 100 (19.5) |
| **Occupation** |  | [21 – 31[ | 126 (24.5) |
| Driver | 12 (2.3) | [31 – 100] | 288 (56) |
| Farmer | 259 (50.4) | **Educational level** |  |
| Student | 110 (21.4) | Illiterate | 50 (9.9) |
| Teacher | 14 (2.7) | Primary | 243 (47.9) |
| Technician | 34 (6.6) | Secondary | 189 (37.3) |
| Trader | 51 (9.9) | Higher | 25 (4.9) |
| Other | 34 (6.6) |  |  |

### Awareness of the targeted NTDs

More than half of respondents (58.4%) indicated that they have never heard about onchocerciasis (Table 2). For those who were aware of this debilitating disease, the most common information sources were community members (42.2%) and health personnel (41.1%). Regarding LF, 379 (73.7%) respondents indicated that they had already heard about this filarial infection, the most common source of information being community members (Table 2). STH infections were well known to the respondents (89.9%), mostly from health personnel (61.5%) (Table 2).

**Table 2.** Knowledge, attitude and practice of populations with regards to onchocerciasis, lymphatic filariasis and soil transmitted helminthiasis

| Indicative questions | Response categories | Oncho<br>N (%) | LF<br>N (%) | STH<br>N (%) |
| --- | --- | --- | --- | --- |
| Have you heard about this NTD? | Yes | 212 (41.2) | 379 (73.7) | 462 (89.9) |
|  | No | 300 (58.4) | 132 (25.7) | 50 (9.7) |
|  | Refuse to answer | 2 (0.4) | 3 (0.6) | 2 (0.4) |
| If « Yes » |  |  |  |  |
| By which means? | Radio | 21 (10.0) | 23 (6.1) | 5 (1.1) |
|  | Television | 5 (2.3) | 11 (2.9) | 4 (0.9) |
|  | Poster | 9 (4.2) | 34 (9.0) | 2 (0.4) |
|  | Health personnel | 86 (41.1) | 111 (29.3) | 284 (61.5) |
|  | Member of the community | 98 (46.2) | 220 (58.0) | 156 (33.8) |
| How can you get this disease? | Flies | 19 (9.0) | 13 (3.4) | NA |
|  | Mosquito | 21 (10.0) | 35 (9.2) | NA |
|  | Inherited from parent/family | 2 (0.9) | 1 (0.3) | NA |
|  | Worms enter through the foot | NA | NA | 4 (0.9) |
|  | Bad hygiene* | NA | NA | 151 (32.7) |
|  | Not washing hands | NA | NA | 22 (4.8) |
|  | Inherited from parent/family | NA | NA | 1 (0.2) |
|  | Don't know | 145 (68.4) | 308 (81.3) | 255 (55.2) |
| What does this disease cause? | Pruritus or itching | 0 (0) | NA | NA |
|  | Skin disease | 6 (2.8) | NA | NA |
|  | Eye disease | 155 (73.1) | NA | NA |
|  | Elephantiasis | NA | 328 (86.5) | NA |
|  | Hydrocele | NA | 9 (2.4) | NA |
|  | Stomachache | NA | NA | 329 (71.2) |
|  | Diarrhea | NA | NA | 75 (16.2) |
|  | Vomiting | NA | NA | 67 (14.5) |
|  | Don't know | 48 (22.6) | 42 (11.1) | 42 (9.1) |
| How can you prevent this disease? | Take Medicine | 41 (19.3) | 57 (15.0) | 136 (29.4) |
|  | Kill flies | 8 (3.8) | 10 (2.6) | NA |
|  | Use a bed net | 8 (3.8) | 5 (1.3) | NA |
|  | Use latrines | NA | NA | 2 (0.4) |
|  | Good Hygiene** | NA | NA | 80 (17.3) |
|  | Don't know | 146 (68.9) | 307 (81.0) | 226 (48.9) |
| What can you do to treat this disease? | Take medicine | 91 (42.9) | 112 (29.6) | 342 (74.0) |
|  | Kill Flies | 0 (0) | 0 (0) | NA |
|  | Use latrines | NA | NA | 2 (0.4) |
|  | Good hygiene** | NA | NA | 2 (0.4) |
|  | Don't know | 112 (52.8) | 234 (61.7) | 112 (24.2) |

### Knowledge of modes of transmission

Among those who were aware of these diseases, only few of them (16.0%) identified black flies as the river blindness transmission agent (Table 2). A similar situation was observed for LF, less than quarter (10%) of the respondents indicating that mosquito act as vector (Table 2). Likewise awareness, respondents’ knowledge of STH transmission was higher as compared to onchocerciasis and LF, though remaining low (Table 2).

### Knowledge about clinical manifestations

Of the respondents who were aware of onchocerciasis, an important proportion of study participants (73.1%) identified eye lesions among clinical signs (Table 2). Elephantiasis of lower limb was reported by 87.0% of those who were aware of LF, while only 2.3% of them identified hydroceles as one of the LF clinical sign (Table 2). Stomachache (71.2%), diarrhea (16.2%) and vomiting (15.0%) were the main clinical manifestations reported by study participants who were aware of STH infections (Table 2).

### Knowledge regarding prevention and treatment approaches

Of those respondents who were aware of onchocerciasis, prevention and treatment approaches were not known by more than half of the respondents. Among those participants who knew that medicines can be used for chemoprevention (19.3%) or chemotherapy (43.1%), only few of them knew that ivermectin was the drug routinely used (Table 2). Overall, 41.0% and 19.8% of the respondents presented with good scores of knowledge and attitudes/practices, respectively (Supplementary material Table S1). Less than quarter (14.2%) of the respondents who were aware of LF choose chemotherapy as the most effective way to prevent this disease, and only few of them indicated that killing mosquitoes (3.0%) by for example using bed nets (1.3%) can be used as another means of LF prevention. As for onchocerciasis, a relatively high proportion of study participants (30.1%) declared that conventional medicines can be used for chemotherapy, but only a few knew that ivermectin is the commonly used drug (Table 3). Overall, 64.4% and 28.8% of the respondents received good scores of knowledge and attitudes/practices, respectively (Supplementary material Table S1). Among respondents who were aware of STH infections, 29.4% and 17.6% reported the use of conventional medicine and applying good hygiene measures, respectively, as the prevention means. Also, a large majority (71.1%) of study participants declared that conventional medicine can be used to treat STH infections, though only a few knew that the routinely used drugs are Ablbendazole and Mebendazole. In addition, 74.0% and 34.6% of the respondents presented with scores of knowledge and attitudes/practices, respectively (Supplementary material Table S1).

### Association between KAP scores of targeted NTDs and different covariates

The univariate analysis has been conducted to assess the association between KAP scores and socio-demographic co-variates (age, level of education, occupation and level of urbanization) recorded in the framework of this study (Supplementary material Table S2). Study participants’ knowledge was positively associated with the level of education (both for onchocerciasis and LF; p < 0.027) and age (LF only; p < 0.001). The respondents’ scores of knowledge about STH were significantly higher amongst teacher and student as compared to the other occupational groups (p < 0.001), better in the Rural settings compared to the urban settings (3.11 vs 2.81, p <0.001). Study participants’ attitudes and practices scores were similar among the different socio-demographic and socio-economic variables for onchocerciasis and lymphatic filariasis, but significantly increased with age (p = 0.031) and were associated with occupation (p < 0.001) for STH.

The multivariate analysis was performed using the CART approach to better identify the socio-demographic and socio-economic determinants associated with KAP of the study participants. Regarding onchocerciasis, the determinants were grouped into 7 classes (Figure 1). Three classes of determinants – class 5 (students or traders over 21 years with at least secondary education and living in urban or semi-urban areas), class 6 (technicians or traders or students over 21 years witch at least secondary education and living in rural areas) and class 7 (teachers) – presented with better knowledge than the reference class 1 (driver, farmer or other trades) (p < 0.006). However, no significant association was found between respondents’ attitudes and practices and these socio-demographic and socio-economic classes (Figure 1, Supplementary material Table S3). As for lymphatic filariasis, CART enable to organize socio-demographic and socio-economic determinants into 8 classes (Figure 2). Study participants from socio-demographic and socio-economic class 8 (drivers or teachers or technician or trader) had better knowledge than individuals from the reference class 7 (farmers or students over 21 years of age living in rural or urban areas) (OR: 2.065, 95% CI: 1.19 – 3.71, p = 0.012). At the contrary of class 8, class 1 (farmers or students under 21 years, having made at most primary) and class 4 (farmers and others over 21 years with a level of education other than the primary and living in semi-urban areas) presented with poor knowledge compared to the reference class 7 (OR: 0.119, 95% CI: 0.03 – 0.31, p <0.001), (OR: 0.290, 95% CI: 0.14 – 0.58, p <0.001), respectively. No significant difference was found between the attitudes and practices scores and the different classes of socio-demographic and socio-economic status of the respondents (p = 0.217) (Supplementary material Table S4). For STH infections, CART led to the organization of socio-demographic and socio-economic determinants into three main classes (Figure 3). Study participants belonging to class 2 (students or other workers who attended) had poor knowledge of STH compared to the reference class 3 (worker driver + farmer + technician + teacher + trader) (OR: 0.359, 95% CI: 0.22 – 0.57, p <0.001). Contrarily to onchocerciasis and lymphatic filariasis, attitudes and practices of study participants with regards to STH infections was associated with socio-demographic and socio-economic determinats (p <0.001). Respondents from class 2 (students or other workers) presented with inapropriate attitudes and practices with regards with STH infections compared to the reference class 3 (worker driver + farmer + technician + teacher + trader) (OR: 0.359, 95% CI: 0.22 – 0.57, p <0.001) (Supplementary material Table S5).

**Fig1.**
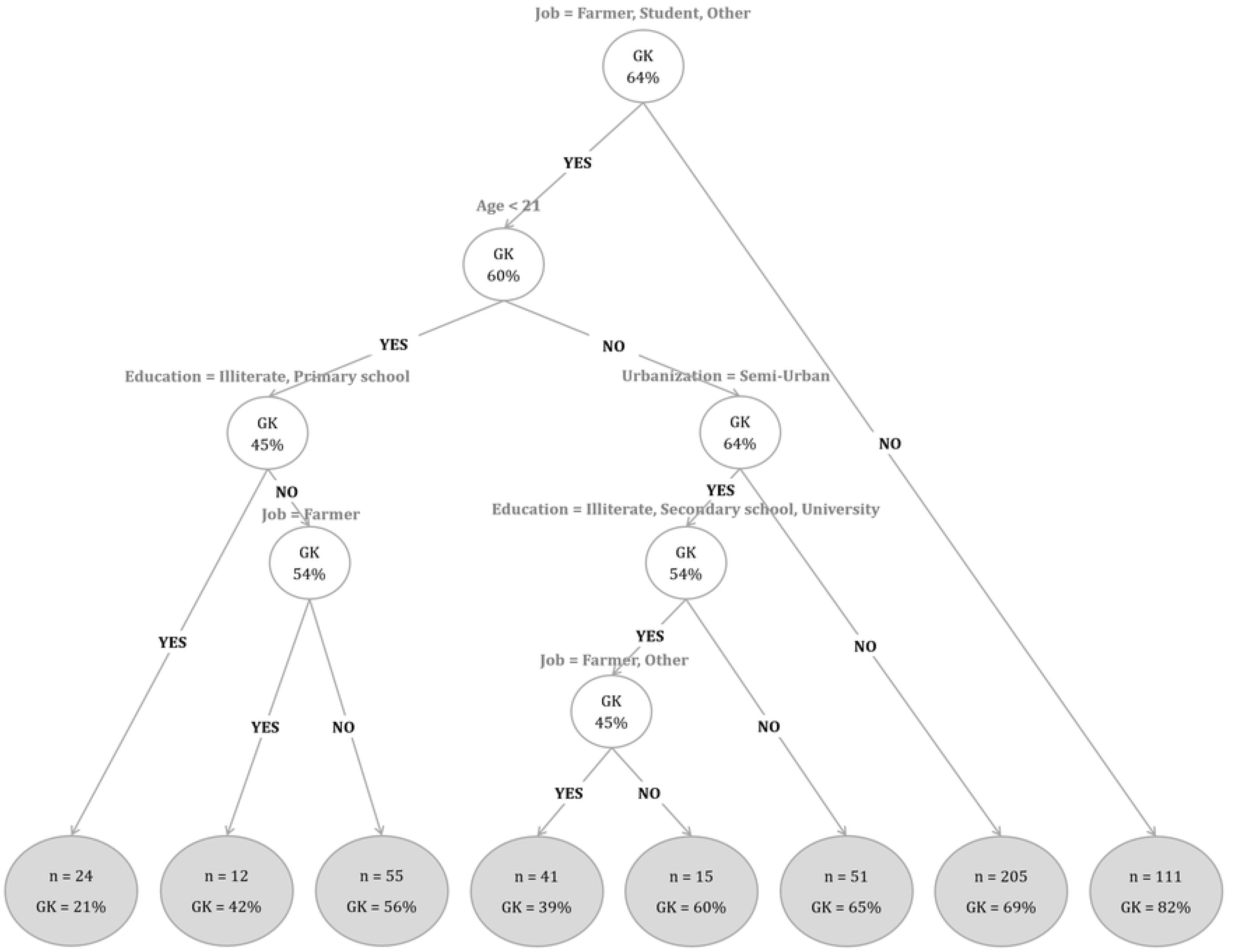
Multivariate analysis using CART to identify determinants of KAP for onchocerciasis. GK indicates good knowledge.

**Fig2.**
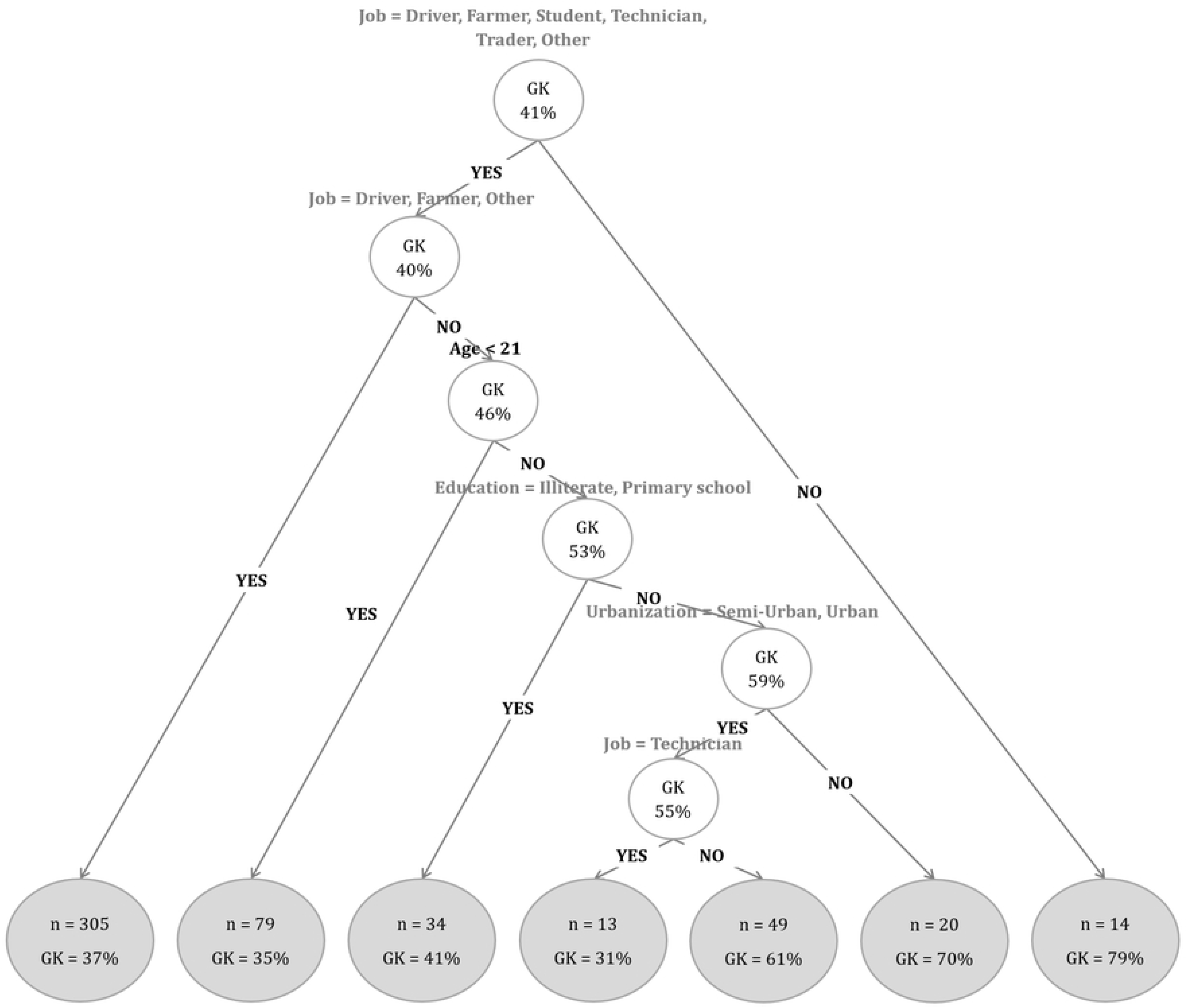
Multivariate analysis using CART to identify determinants of KAP for LF. GK indicates good knowledge.

**Fig3.**
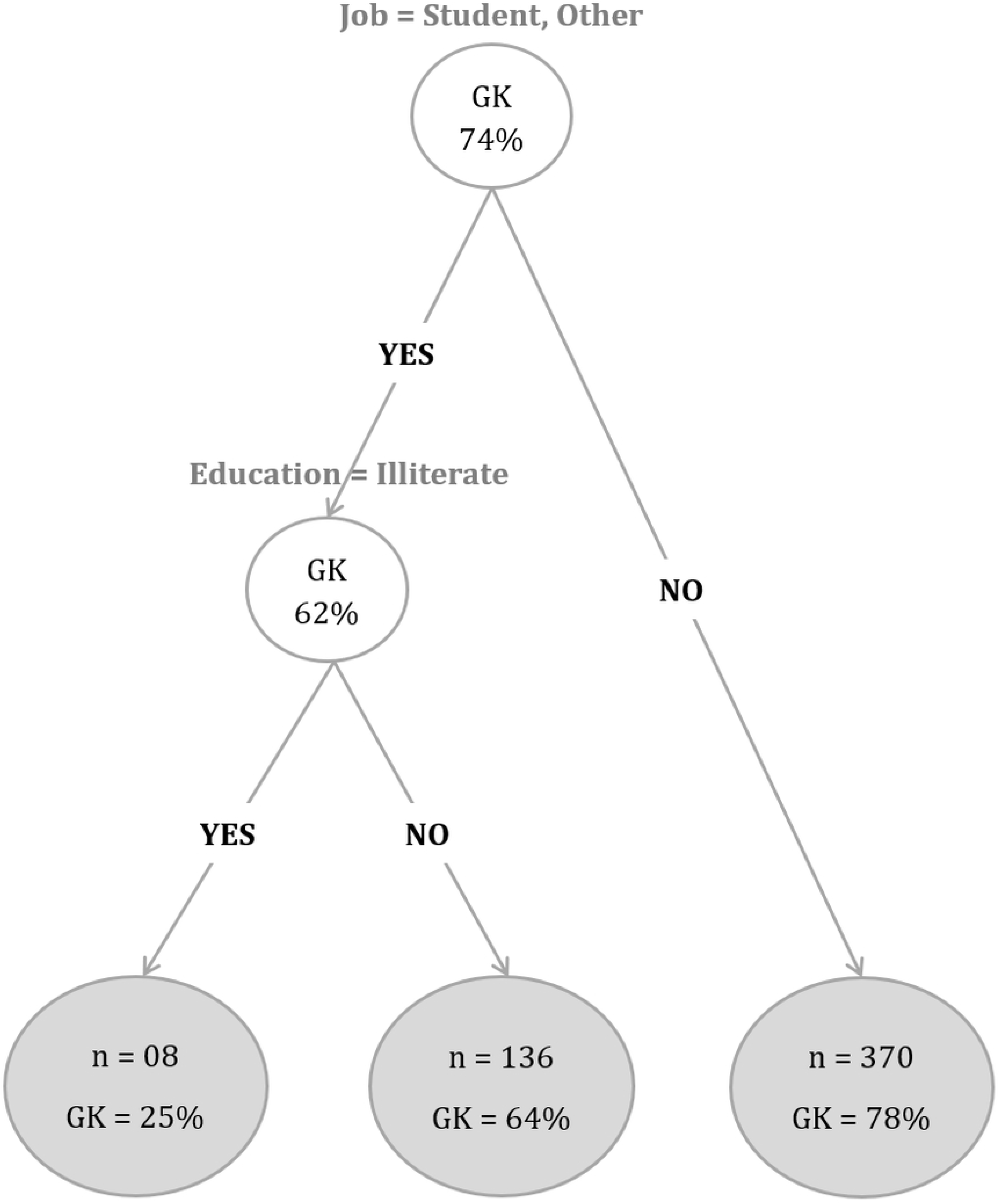
Multivariate analysis using CART to identify determinants of KAP for STH. GK indicates good knowledge.

### Relationship between knowledge of respondents and their attitudes/practices

Onchocerciasis knowledge was significantly associated with respondents’ attitudes and practices (Correlation Coefficient (rho) = 0.64, p <0.001), study participants with good knowledge exhibiting the better attitudes and practices. A similar positive relationship trends between respondents’ knowledge about LF and STH infections, and their attitudes and practices was also observed (rho = 0.46, p <0.001) and (rho = 0.46, p <0.001), respectively.

## Discussion

The purpose of this study was to assess, through an integrated approach, the attitudes and practices of the Mbengwi health district populations, in relation to their level of knowledge regarding three of the five major NTDs endemic in this area.

The awareness of study participants was quite variable from one NTD to another. Indeed, more than half of the study participants declared that they had never heard about onchocerciasis, though the fight against this disease is ongoing and already implemented in this health district for more than 15 years. These findings are contrasting to those collected in Ethiopia [19] were the majority of the study participants were familiar with onchocerciasis; this might be explained by the fact that this filarial disease in not as severe in the Mbengwi health district () as in Ethiopia where the endemicity of the disease was high, ranging from 6.9% in the Quara District in northwest Ethiopia, to 85.3% in Teppi in South western [20]. Unlike onchocerciasis, the majority of participants (73.7%) had already heard about LF, from a community member for most of them. This can be explained by the fact that the disease is commonly called elephantiasis, in reference to its most important and visible clinical manifestation. However, these findings are not in line with those collected in Nigeria which revealed that although the region was endemic to LF, the majority of participants (82.1%) were not aware of the disease [21]. Likewise lymphatic filariasis, most of the respondents (89.9%) were well aware of STH infections, mostly from health personnel (61.5%). Indeed, STH are the most widespread NTD over the world, with more than 1.5 billion people infected (~1/4 of the world population). These results corroborate those by Yusof et al. (2017) [22] and Nath et al. (2018) [23] who reported high awareness about STH (62.5%), health workers being the main source of information.

The knowledge of routes of transmission was in general poor, regardless the targeted disease. Indeed, among those who have ever heard about onchocerciasis, 68.3% didn’t knew that the vector is black fly, likely due to the low abundance of this vector in the study area. These results are not in line with the findings by Yirga et al. (2008) [24] in a study conducted in Southwest Ethiopia where the majority of the respondents associated the disease to the bite of black flies. Likewise onchocerciasis, the majority of participants did not know that mosquito is the vector of lymphatic filariasis, as was also previously observed elsewhere [25–26]. Regarding STH infections, 55.2% of participants didn’t know how this NTD is transmitted, contrarily to the finding by Nath et al. (2018) [23] where school-aged children were aware that STH are transmitted through contaminated soil (58.1%) and unhygienic practices (55.7%). In Cameroon, the Schistosomiasis and Soil Transmitted Helminthiasis National Control program is based on the deworming of school-aged children by their teachers. Thus, during the treatment campaigns in primary schools, the out-target population (included in this study) is not educated about the disease and might explain why study participants were so little familiar with the modes of transmission of STH.

Regarding knowledge of clinical manifestations, most of the study participants (73.1%) identified eye disease as a clinical manifestation of onchocerciasis, probably in reference to the common name of onchocerciasis, river blindness. Such high level of clinical manifestation knowledge has been observed in a study by Weldegebreal et al. (2014) [19], the most reported clinical sign being however itching. A similar trend was observed with lymphatic filariasis, 87.0% of respondents reporting elephantiasis as the main clinical manifestation. These results corroborate those by Amaechi and colleagues (2016) [21] who reported swellings (84.6%) as the most common responses symptoms in Omi irrigation community at North Central of Nigeria. Also, most of the study participants identified stomachache and diarrhea (87.4%) as the main STH clinical manifestations, as was previously reported in a study conducted in two rural communities of western Ivory Coast [27]. In general, the knowledge of populations as regards with clinical signs of any of the targeted NTDs was high, more likely because these diseases are usually assimilated to their most prominent symptoms.

The majority of study participants (69.1%) was not aware of onchocerciasis prevention means, and almost half of them (53.0%) ignored how to treat this filarial disease. This was quite surprising, especially because the fight against this disease through the CDTI is implemented in this health district since more than 15 years, with acceptable therapeutic coverage. It is worth to however mention that CDTI is critically deviating from its initial pathway, with absence of health education at the community level, reduced commitment of CDDs, poor training and supervision, no more self-monitoring by communities …, and might likely explain why populations can adhere to treatment without knowing onchocerciasis prevention and control measures. These results are not in line with those of Weldegebreal et al. (2014) [19] in Nigeria where almost all (93.3%) the respondents believed that onchocerciasis is preventable, and 88.4% of them knew that ivermectin is the drug of choice. Regarding lymphatic filariasis, a very little proportion knew LF prevention and treatment strategies, contrarily to what was previously reported in the Omi community in Nigeria, where 61.8% knew how to prevent lymphatic filariasis, and 49.8% reported the use of anti-filarial drug [21]. This might be explained by the deviation observed in CDTI implementation, onchocerciasis and lymphatic filariasis being implemented following the same approach and by the same actors. More than half of the study participants knew how to prevent STH infections, and 71.1% of them knew that this disease can be treated using drugs, though almost none knew exactly which drug is usually used. These findings were in line with those by Nath and colleagues (2018) [23] in Bangladesh where 64.4% of school-aged children interviewed declared preventing STH by washing hands after defecation, and 75.6% of them knew that to control STH they should take drugs.

Multivariate logistic regression revealed that knowledge of these three group of diseases was associated with age of enrollees, their level of education, their occupation and the level of urbanization. Indeed, those individuals aged ≥21 years old, who have attended at least the secondary school, who were student/teachers and living in urban settings exhibited in general better knowledge as regards to these diseases. Though these factors have also been observed in previous onchocerciasis, lymphatic filariasis and STH related studies [21, 28]), other determinants, not necessarily evaluated in the framework of this study, might also be associated with of knowledge of populations. For example, in a previous study on onchocerciasis [19], only ethnicity was associated with knowledge, attitudes and practices of populations.

Finally, a positive association was found between knowledge of respondents and their attitudes and practices with regard to the three targeted NTDs, suggesting that the better populations are educated, the most they can be aware of the importance of control measures and thus better comply.

This study revealed that study participants exhibit poor knowledge, attitudes and practices as regards to these three PCT-based highly prevalent diseases, although they are endemic in the study area and MDA administered yearly since decades. Misconceptions can seriously affect the adherence and contribution of populations to the success of PCTs, and consequently elimination of targeted diseases. Since lack/poor knowledge and wrong beliefs could lead to inappropriate control measures, it appears compulsory to improve individual knowledge, with a focus on the importance and rationale behind MDA, to optimize their attitudes and practices, especially community participation to PCTs.

## Acknowledgements

The authors are grateful to the population of the Mbengwi health district who willingly accept to participate in this study.

## Supporting information

**S1 Table. Descriptive statistics of scores of KAP (DOCX)**

**S2 Table. Univariate analysis of NTDs by socio-demographic characteristics (DOCX)**

**S3. Table. Analysis of behavioral classes and determinants of KAP about onchocerciasis (DOCX)**

**S4. Table. Analysis of behavioral classes and determinants of KAP about FL (DOCX)**

**S5. Table. Analysis of behavioral classes and determinants of KAP about STH (DOCX)**

